# Acute toxicity effects of pesticides on predatory snout mites (family Bdellidae)

**DOI:** 10.1101/2023.10.17.562818

**Authors:** Rosemary A. Knapp, Luis Mata, Robert McDougall, Qiong Yang, Ary A. Hoffmann, Paul A. Umina

**Author notes:** Corresponding author: Rosemary A. Knapp.

## Abstract

Predatory mites biologically control a range of arthropod crop pests and are often central to agricultural IPM strategies globally. Conflict between chemical and biological pest control has prompted increasing interest in selective pesticides with fewer off-target impacts on beneficial invertebrates, including predatory mites. However, the range of predatory mite species included in standardised pesticide toxicity assessments does not match the diversity of naturally-occurring species contributing to biocontrol, with most testing carried out on species from the family Phytoseiidae. Here, we aim to bridge this knowledge gap by investigating the impacts of 22 agricultural pesticides on the predatory snout mite *Odontoscirus lapidaria* (Kramer) (family Bdellidae). Using internationally standardised testing methodologies, we identified several active ingredients with minimal impact on *O. lapidaria* mortality, including *Bacillus thuringiensis*, nuclear polyhedrosis virus, flonicamid, afidopyropen, chlorantraniliprole and cyantraniliprole, which may therefore be good candidates for IPM strategies utilising both chemical and biological control. Importantly, we reveal differences between Bdellidae and Phytoseiidae in responses to a number of chemicals, including the miticides diafenthiuron and abamectin, highlighting the risk of making generalisations around acute toxicity based on tests with one beneficial mite family. We also explored the impacts of several pesticides on a second Bdellidae species and found differences in the response to chlorpyrifos compared with *O. lapidaria*, further highlighting the taxon-specific nature of non-target toxicity effects.

## 1. Introduction

Protecting field crops from damage caused by arthropod pests is essential to ensure food quality and yield requirements are met globally (Oerke, 2006). While chemical control remains the major tool to defend against crop pests, the threat of pesticide resistance and increasing focus on environmental sustainability has driven a widespread interest in reducing their use through integrated pest management (IPM) strategies (Hoffmann et al., 2008; Phillips et al., 1989). Biological control from natural enemies, such as predators and parasitoids, is central to IPM and is an important line of defence against a variety of globally important arthropod pests (Altieri, 1999; Power, 2010).

Predatory mites play a key role as biological control agents in many agricultural environments (Knapp et al., 2018). These generalist predators can provide sustained control of a range of crop pests (Carrillo et al., 2015; Tixier, 2018), reducing or removing the need for pesticide applications in some situations (Blommers, 1994; Prischmann et al., 2006). Phytoseiid mites (family Phytoseiidae), including species in genera *Amblyseius*, *Euseius*, *Neoseiulus* and *Phytoseiulus*, are important predators of pest spider mites such as *Tetranychus urticae*, *Panonychus citri* and *Panonychus ulmi* (Fountain and Medd, 2015; Fraulo and Liburd, 2007; Van Leeuwen et al., 2015). Deliberate introductions of phytoseiid mites to crops through augmentative release programs are common. For instance, *Neoseiulus californicus* is routinely introduced to control the two-spotted spider mite *T. urticae* (Fraulo and Liburd 2007). Though less common, soil-dwelling mites (family Laelapidae) are also commercially available for biological control. For example, A*ndrolaelaps casalis*, *Gaeolaelaps aculeifer* and *Stratiolaelaps spp.* are used to control pests such as the Western flower thrip *Frankliniella occidentalis* and dark-winged fungus gnats (Diptera: Sciaridae) (Moreira and de Moraes, 2015).

Importantly, both crop and non-crop vegetation within agroecosystems host a range of naturally-occurring predatory mite species, beyond those potentially used in augmentative release programs (Tixier, 2018). Pest suppression from resident predatory mites can be heightened by landscape and habitat manipulation management actions aimed at maintaining and promoting beneficial mite diversity (Etienne et al., 2021; Möth et al., 2021). Another conservation biological control strategy is the use of selective pesticides, which have been specifically designed to reduce their off-target impacts on predatory mites and other beneficial species, over non-selective broad-spectrums (Holloway et al., 2008). For instance, pesticide application regimes favouring selective active ingredients such as *Bacillus thuringiensis (Bt)* over synthetic pyrethroids (SPs) and organophosphates (OPs) are compatible with biological control of the pest mite *Panonychus ulmi* using predatory mites in apple orchards (Agnello et al., 2003). In this respect, maintaining the diversity of naturally-occurring predatory mites to aid biological control relies upon a thorough understanding of the off-target impacts of pesticides on a range of species (Duso et al., 2020).

Knowledge of the impacts of pesticides on predatory mites is currently limited by a lack of diversity in the species typically studied (Overton et al., 2021). Most research has focused on phytoseiid mites, constituting almost 90% of papers using the keyword ‘predatory mites’ in Scopus (Duso et al., 2020). Standardised pesticide toxicity assessments typically do not extend beyond a few representative commercially available Phytoseiid mite species, such as *Phytoseiulus persimilis*, *Neoseiulus spp.*, *Typhlodromus pyri* and *Amblyseius spp.* (Biobest, 2019; Bostanian, 1985, 2006; Choi, 2003; Cote, 2002; Cuthberson, 2012; Duso, 1992; Kaplan et al., 2012; Kim, 2005; Kishimoto, 2018; Zhang, 1990), with some of these acting as ‘indicator’ species in ecotoxicological studies informing pesticide regulations (Candolfi et al., 1999). However, it is unclear how representative phytoseiid mites are of the broader diversity of predatory mite families contributing to biological control, and few studies have investigated the toxicity of agricultural pesticides towards non-phytoseiid predatory mites.

Snout mites (family Bdellidae) are a particularly understudied group of predatory mites, despite being globally distributed (Hernandes et al., 2016), among the first mites ever described (Linnaeus, 1758), and one of the earliest mite groups observed undertaking pest control (Womersley, 1933). The pasture snout mite, *Odontoscirus lapidaria*, has in particular been observed predating on pests such as the lucerne flea, *Sminthurus viridis*, in Australia (Currie, 1934; Wallace, 1967; Womersley, 1933), prompting later introductions to South Africa for biological control (Wallace and Walters, 1974). More recently, the spiny snout mite, *Neomulgus capillatu*s, has been observed in a suspected predator-prey relationship with another important agricultural pest, the redlegged earth mite, *Halotydeus destructor* (Bell and Willoughby, 2003).

Other than a single study on the impacts of SPs and OPs on *O. lapidaria* (Roberts et al., 2011), the majority of currently available agricultural pesticides have not been tested on Bdellidae mites. In particular, it is unclear whether bdellids have similar responses to pesticides deemed to be ‘selective’ through previous testing on phytoseiid species. This work aims to help bridge this gap by testing a range of pesticides on two bdellid species to diversify the predatory mite families from which IPM recommendations can be drawn. Specifically, we address the following research questions (RQs):

RQ1 – what are acute toxicities of 22 common agricultural pesticides towards *O. lapidaria*?

RQ2 – what are the temporal impacts of pesticides at multiple application rates?

RQ3 – how comparable are *O. lapidaria’s* responses to a sub-set of pesticides with a second distinct bdellid species?

## 2. Methods

### 2.1. Collection of snout mites

Snout mites (family Bdellidae) were collected from two sites in Victoria, Australia, during the winters of 2021 – 2023. Site A (Hurstbridge, 37°38’04”S 145°12’21”E) was a natural grassland with no recorded pesticide applications within the past 10 years. Site B (Inverleigh, 38°10’37”S, 144°00’10”E) was a shelterbelt located between two fields used for rotational grain cropping. The shelterbelt was approximately 15 m wide with an open canopy of native eucalypt trees and an understory that was a mix of native grasses and exotic herbaceous plants with no record of direct pesticide application.

Collectionsweremadebysuctionsampling using a blower vacuum (Stihl BG86, Germany) modified with a metal sieve fitted to the vacuum spout. This method is commonly used to sample ground-dwelling invertebrates and has minimal impact on their condition (e.g., Roberts et al., 2009, 2011; Umina and Hoffmann, 2005). All collected invertebrates were transported back to the laboratory in ventilated plastic containers with paper towel and vegetation to maintain humidity. Snout mites were then separated using a handheld suction device or paintbrush and stored at 4 °C for a maximum of two days before acute toxicity bioassays.

### 2.2. DNA barcoding

As the taxonomy of Australian Bdellidae is poorly developed and it is difficult to distinguish taxa through morphological examinations, a sub-sample of snout mites (recognized by their elongated gnathosoma and terminal setae) from each field collection (3-10) were sequenced using mitochondrial COI as a DNA barcode to identify species from each site. DNA was extracted using 150 μL 5% Chelex 100 resin (Bio-Rad Laboratories, Hercules, United States) according to methods described previously (Lee et al 2012). The universal barcode region of the CO1 gene (658 bp) of individual mites was amplified using the generic primers (LCO1490 5’ GGTCAACAAATCATAAAGATATTGG 3’ and HCO2198 5’ TAAACTTCAGGGTGACCAAAAAATCA 3’) (Folmer et al 1994). PCR amplifications were performed in a Thermal Cycler (Eppendorf, Hamburg, Germany) with an adjusted annealing temperature of 56 °C. Amplicons were sequenced via Sanger Sequencing (Macrogen, Seoul, South Korea). Sequences were matched to the DNA barcode reference library on BOLD to confirm taxonomy identifications.

All mite sequences from Site A were found to match the pasture snout mite, *O. lapidaria* (BOLD: ADX6309, 100% similarity). Site B mites could not be identified beyond family level (Bdellidae, average 89% similarity), but comparison of DNA sequences shows that they are a different species to *O. lapidaria*, as sequence similarity between them was only 69.4% (Figure S1). These are henceforth referred to as ‘Bdellidae sp. B’.

### 2.3. Pesticides

A comprehensive range of pesticide formulations was tested, comprising 14 Mode of Action (MoA) groups and 22 unique active ingredients (Table 1). Pesticides were tested at two rates (high and low) based on the Maximum Registered Field Rate (MRFR) of their active ingredients in Australian grain crops (APVMA, 2021). High rates are 100% of the MRFR and low rates either 10% of the MRFR or the Lowest Registered Field Rate (LRFR) if an active ingredient was registered across a particularly wide range of rates (e.g. pirimicarb, chlorpyrifos and dimethoate). Solutions of each pesticide rate were made up by diluting the necessary volume of product in deionised water.

**Table 1 –.**
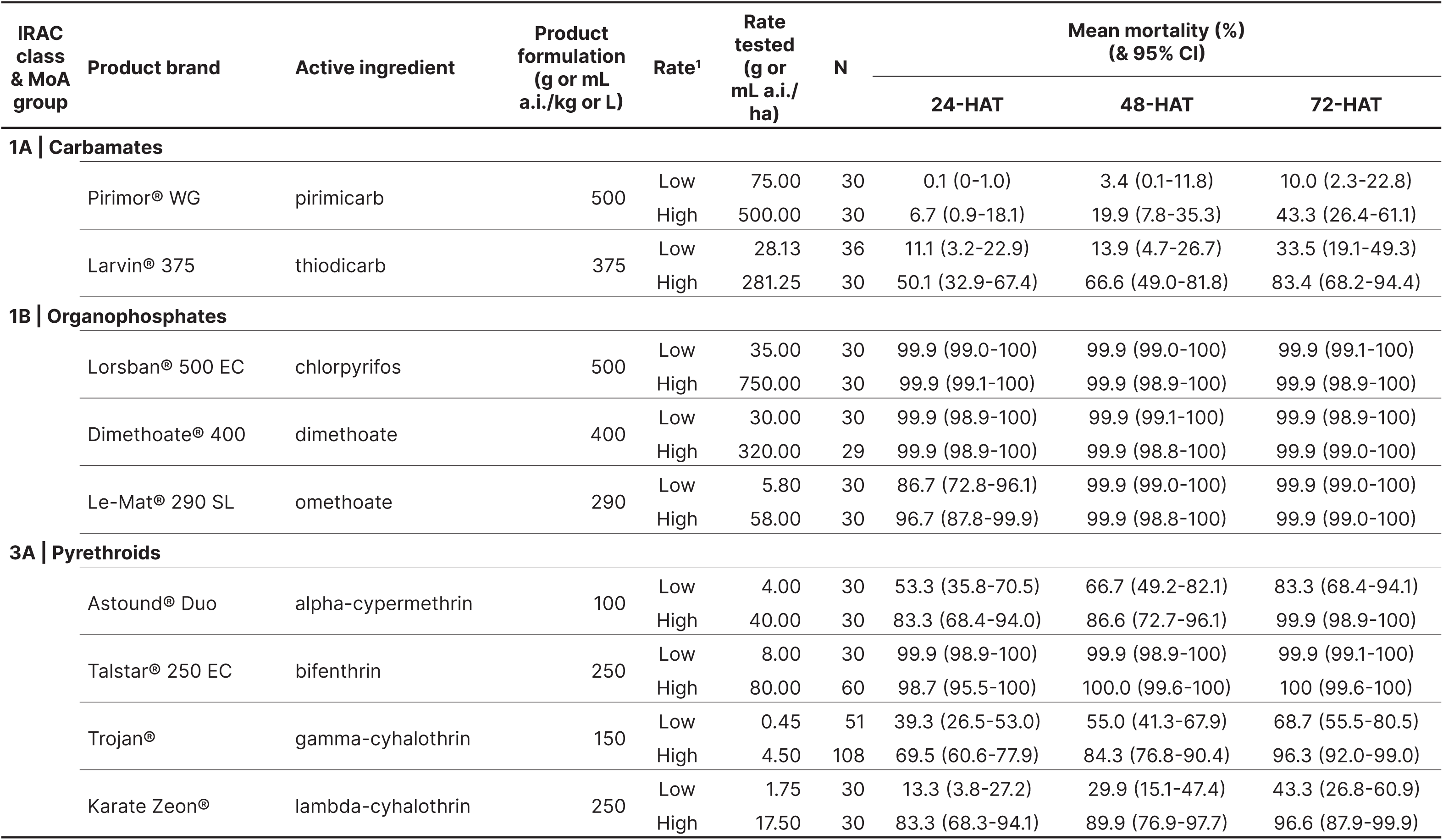
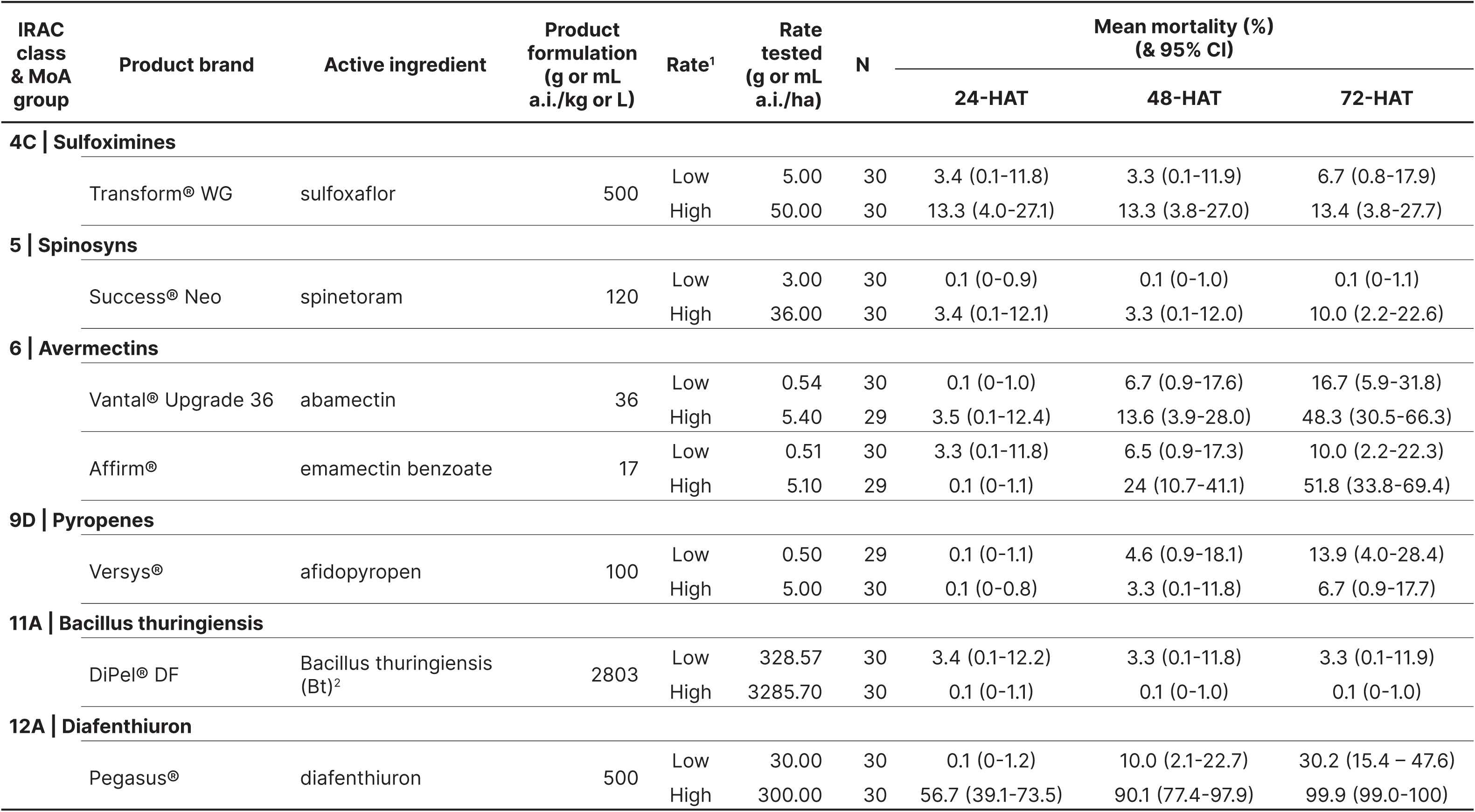
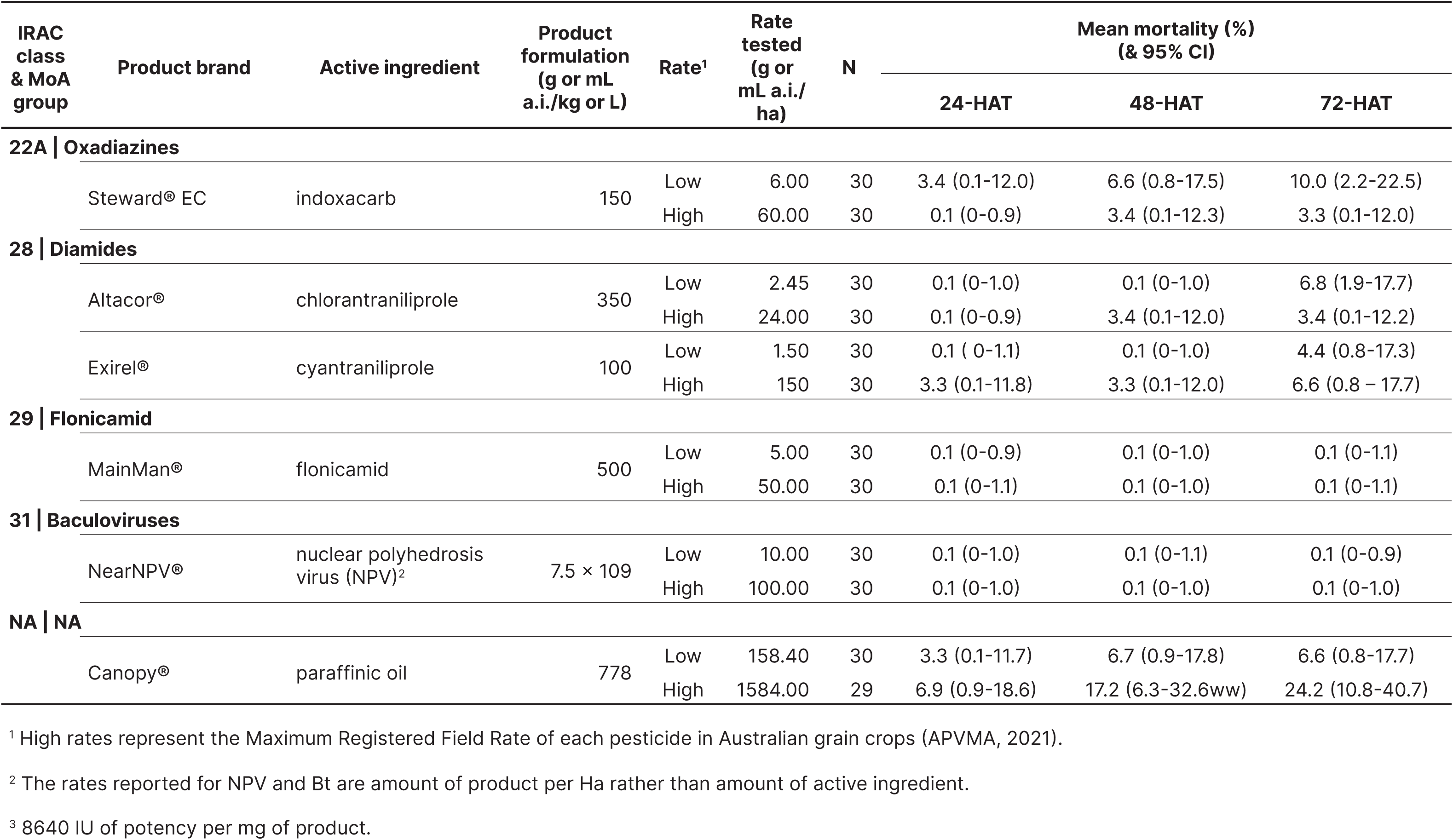
Posterior estimates for *O. lapidaria* mortality (expressed as a percentage) to the 22 tested pesticides at the high and low rates after 24, 48 and 72-HAT expsoure, including details on Insecticide Resistance Action Committee (IRAC) classification, Mode of Action (MoA), product brand and formulation, rate tested, and numbers of mites tested (N). Gamma-cyhalothrin and bifenthrin were used as positive controls across multiple bioassays, hence N > 30. Cases where N < 30 represent instances of mite escapees or insufficient mite numbers. Comprehensive information on the treatments tested in each bioassay and associated negative control mortalities are given in Table S1. HAT: hours after treatment; CI: credible interval.

### 2.4. Acute toxicity bioassays

The acute toxicity of all 22 pesticides at both rates was tested on *O. lapidaria* through laboratory bioassays (RQ1 and RQ2). For the cross-species comparison (RQ3), five pesticides representing a range of MoA groups and acute toxicities (abamectin, chlorpyrifos, diafenthiuron, paraffinic oil and spinetoram) were also tested on Bdellidae sp. B at the high rates only.

The large number of treatments in this study (49 in total) required data to be collected over multiple discrete bioassays. The same methodology was used across all bioassays, with each including a negative control (deionised water) and positive control (typically gamma-cyhalothrin or bifenthrin) to ensure consistency in mite responses.

In all bioassays, snout mites were exposed to dried pesticide spray residues in line with International Organisation for Biological Control (IOBC) protocols (Sterk et al., 1999) and previous work (Overton et al., 2023). A Potter Spray Tower (Burkard Manufacturing, Rickmansworth, United Kingdom) was used to coat 35 mm petri dish bases with even and consistent spray deposits of 1 mg/cm^2^, equivalent to a field application rate of 100 L/ha of diluted pesticide. Three dishes were sprayed at once, and a total of 30 replicates (where one replicate = one dish) were carried out for each treatment unless otherwise stated (Table 1). Residues were allowed to dry in a fume hood for a minimum of 30 min after spraying.

Once dry, a single mite was transferred into each petri dish along with a vetch leaf (*Vicia sativa* cv. Blanchefleur) to maintain humidity. Dishes were sealed with parafilm to prevent mites escaping. To prevent desiccation and reduce possible fumigant effects between pesticide treatments, all dishes within a treatment were placed in 85 L sealed storage containers (Inadox, Melbourne, Australia) with a piece of damp paper towel and kept at room temperature. Individuals were scored as alive (actively moving), dead (no movement) or incapacitated (inhibited movement) at three timepoints: 24, 48 and 72 h after treatment (HAT). Incapacitated individuals were pooled with dead individuals for analysis, as they are unlikely to survive or exhibit normal predatory behaviours in the wild (Hoffmann et al., 1997; Roberts et al., 2011).

### 2.5 Statistical analysis

A series of generalised linear models were used to assess mortality (Kéry, 2010), which were specified as:

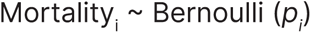

where *p_i_* is the probability that the tested mite in replicate petri dish *i* will be dead or incapacitated. In all models, the linear predictors were specified on the logit-probability scale, with the logit link function defined as ln(p/1-p).

The linear predictor ofthe model assessing how *O. lapidaria* mortality varied across chemicals (RQ1) was specified as:

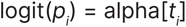

where *t_i_* is an integer indexing the 22 chemical treatments and alpha the treatment-level fix effects. This model only included the high rate and 48-HAT data, in order to make comparisons with previous studies benchmarked against current IOBC standards (Roberts et al., 1999), as well as ongoing work on a wider range of beneficial invertebrates (Mata et al., unpubl. data; Overton et al., 2023; Umina et al., 2023).

The linear predictors of the models assessing how *O. lapidaria* mortality varied across chemicals and HATs (RQ2) were specified as:

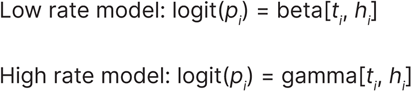

where *t_i_* is an integer indexing the 22 chemical treatments; *h_i_* an integer indexing the three HAT timepoints; and beta and gamma the combined chemical/ HAT treatment-level fix effects, which were specified as two-dimensional matrices. Pesticide rates were modelled in separate batches for simplicity; an alternative approach that would have yielded the same estimates would have been to specify the fix-effects as a three-dimensional matrix with a corresponding integer indexing the two rates.

The linear predictor of the model assessing how *O. lapidaria* and Bdellidae sp. B mortality varied across chemicals (RQ3) was specified as:

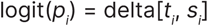

where *t_i_* is an integer indexing the five chemical treatments; *s_i_* an integer indexing the two species; and delta the combined chemical/species treatment-level fix effects, which were specified as a two-dimensional matrix. As with the *O. lapidaria* mortality across chemical analysis (RQ1), this model only included the high pesticide rate and 48-HAT data.

Abbott’s corrections were not used in any analyses, as background (negative) control mortality was consistently under 10% across all bioassays (Table S1).

All model parameters were estimated under Bayesian inference, using Markov Chain Monte Carlo (MCMC) simulations to draw samples from the parameters’ posterior distributions. All fix effect parameters were assigned to non-informative Normal priors (mean=0, sd=32). Models were implemented in JAGS (Plummer, 2003), as accessed through the R package jagsUI (Kellner, 2016). We used three chains of 5,000 iterations, discarding the first 500 in each chain as burn-in. MCMC chains were visually inspected and the values of the Gelman-Rubin statistic used to verify acceptable convergence levels of R-hat < 1.1 (Gelman and Hill, 2007).

Two post-analysis steps were taken to assist with the interpretation of modelling outputs. Firstly, the posterior estimates from the *O. lapidaria* models (RQ1 and RQ2) were used to map the probability or probabilities that the mortality response of *O. lapidaria* to a given chemical (RQ1) at a given rate and HAT (RQ2) will be classified into one or more of the IOBC toxicity categories (Hassan, 1992; Sterk et al., 1999), where 1 = ‘harmless’ (<30% mortality), 2 = ‘slightly harmful’ (30-79% mortality), 3 = ‘moderately harmful’ (80-99% mortality), and 4 = ‘harmful’ (>99% mortality). Secondly, the posterior estimates from all models were run through an algorithm designed to generate group effects triangular tables. The algorithm first compared the 95% credible intervals (CI) of each group pair (e.g. mortalities of chemical A and B in RQ1), either estimating the mean group effect (i.e. the multiplicative effect separating the smaller effect from the larger) when the group pair’s CIs do not overlap or assigning a ‘not statistically different’ (nsd) tag when the CIs overlapped.

The mean group effect (MGE) was calculated as:

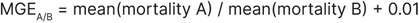

where

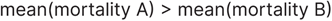

MGE was mathematically constrained to be bounded between 0 and 100 by limiting the number of decimal points used in the calculations to three. 0.01 was added to the smaller effect, which prevents the MGE estimation from diverging towards infinity in cases where the mean mortality was zero.

## 3. Results

### 3.1. Standardised acute toxicity (RQ1)

The acute toxicity response of *O. lapidaria* varied substantially across the 22 tested pesticides (Figure 1, Table 1). When the results for 48-HAT at the high rate are categorised according to IOBC classifications (Sterk et al., 1999), every OP tested (i.e. chlorpyrifos, dimethoate and omethoate) and one SP (bifenthrin) was classified as ‘harmful’, causing on average > 99% mortality (Figure 1, Table 1). Conversely, *Bt* and nuclear polyhedrosis virus (NPV), both biological insecticides, caused < 1% mortality on average and were classified as ‘harmless’ (Figure 1, Table 1). Flonicamid, afidopyropen, chlorantraniliprole, cyantraniliprole, indoxacarb, spinetoram, sulfoxaflor, abamectin, paraffinic oil, pirimicarb, and emamectin benzoate were also classified as ‘harmless’ to *O. lapidaria* (<30% mortality; Figure 1, Table 1). Thiodicarb was the only active classed as ‘slightly harmful’ (30-79% mortality; Figure 1, Table 1), while diafenthiuron and all SPs tested aside from bifenthrin (i.e. gamma-cyhalothrin, lambda-cyhalothrin, alpha-cypermethrin) were classified as ‘moderately harmful’ to *O. lapidaria* (80-99% mortality; Figure 1, Table 1).

**Figure 1 –.**
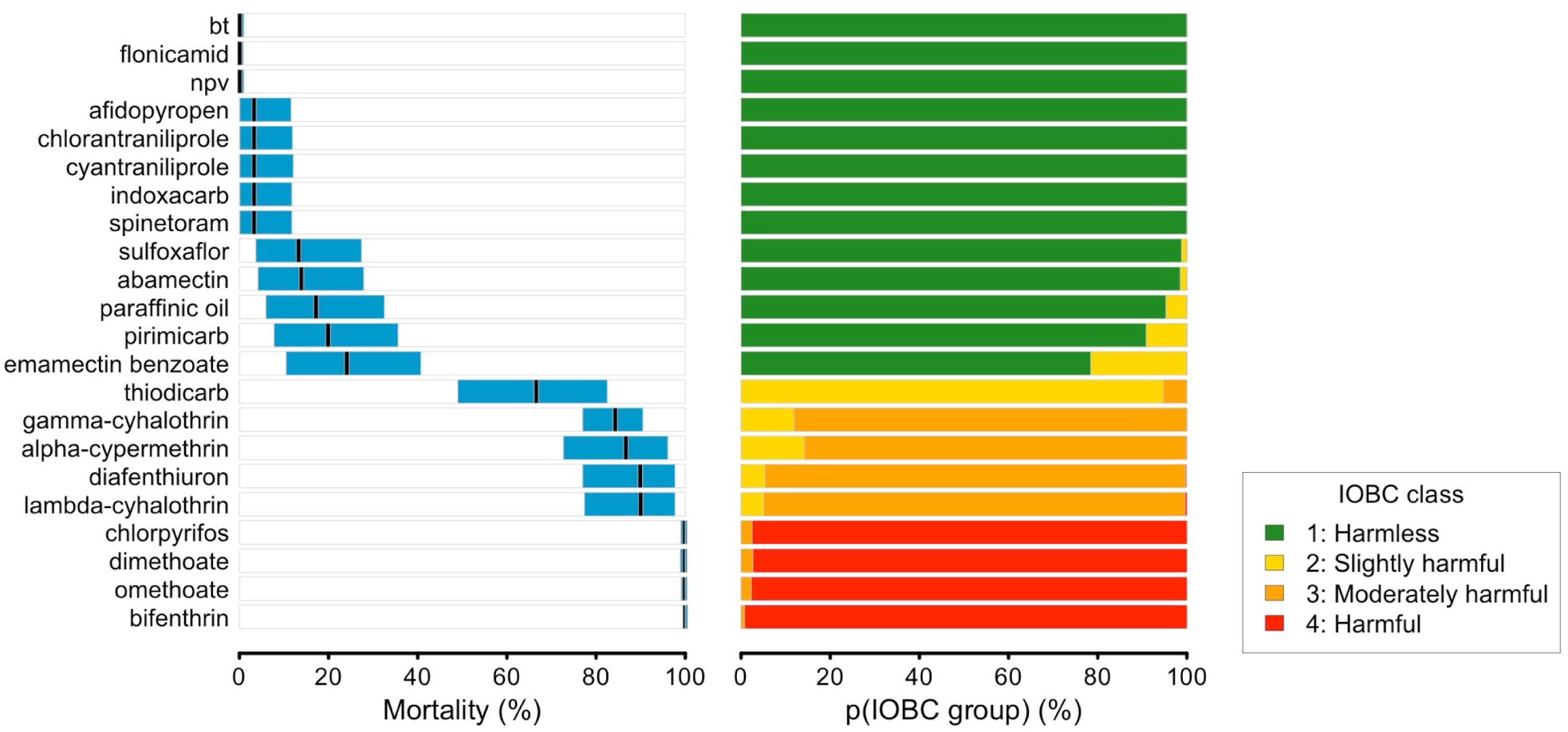
*O. lapidaria* mortality following exposure to high rates of pesticide residues at 48-HAT. Pesticide rates are as described in Table 1. Black lines indicate the mean response and the blue rectangles the associated statistical uncertainty (95% CI). Non-overlapping rectangles denote statistical differences. Pairwise statistical comparisons and mean group effects are given in Table S2. The right-hand panel shows the probability or probabilities that the mortality response of *O. lapidaria* to the given chemical treatment will be categorised into one or more IOBC toxicity classifications, where 1 = ‘harmless’ (< 30% mortality; green), 2 = ‘slightly harmful’ (30-79% mortality; yellow), 3 = ‘moderately harmful’ (80-99% mortality; orange), and 4 = ‘harmful’ (> 99% mortality, red). p: probability; IOBC: International Organisation for Biological Control; HAT: hours after treatment; CI: credible interval.

In general, *O. lapidaria* responded similarly to pesticides whose active ingredients shared the same MoA. At high rates and 48-HAT, there were no statistical within-MoA group differences in mortality for all OPs (Group 1B; chlorpyrifos, dimethoate and omethoate), both avermectins (Group 6; abamectin and emamectin benzoate), both diamides (Group 28; chlorantraniliprole and cyantraniliprole), and for the majority of SPs (Group 3A; alpha-cypermethrin, gamma-cyhalothrin and lambda-cyhalothrin; Table S2). However, bifenthrin was more lethal than the other SPs, causing on average 1.17, 1.14 and 1.10 times higher mortality than gamma-cyhalothrin, alpha-cypermethrin and lambda-cyhalothrin, respectively (Table S2). Further, thiodicarb (classified as ‘slightly harmful’; average 66.7% mortality) was average 3.34 times more lethal than pirimicarb (classified as ‘harmless’; on average 20% mortality), despite both actives being carbamates (Group 1A; Table S2).

### 3.2. Differences in acute toxicity with rate and exposure time (RQ2)

High pesticide rates caused either the same or higher *O. lapidaria* mortality than low rates for all active ingredients tested, and an increased probability of falling into a higher IOBC classification (Figure 2). This effect was most pronounced for mid-range active ingredients classified as ‘slightly harmful’ and ‘moderately harmful’. For instance, at 48-HAT, lambda-cyhalothrin showed strong rate-specific effects on *O. lapidaria* mortality, increasing from an average of 29.9% when exposed to the low rate to 89.9% when exposed to the high rate (Table 1, Figure 2). This corresponds with an increase in the probability of being classified as ‘moderately harmful’ from 0% to 94.4% (Figure 2; Table S3). Conversely, rate had little effect on the toxicity of OPs, all of which caused on average >99% mortality by 48-HAT and had >97% probability of being classed as ‘harmful’ to *O. lapidaria* even at the low rates tested (Figure 2; Table S3).

**Figure 2 –.**
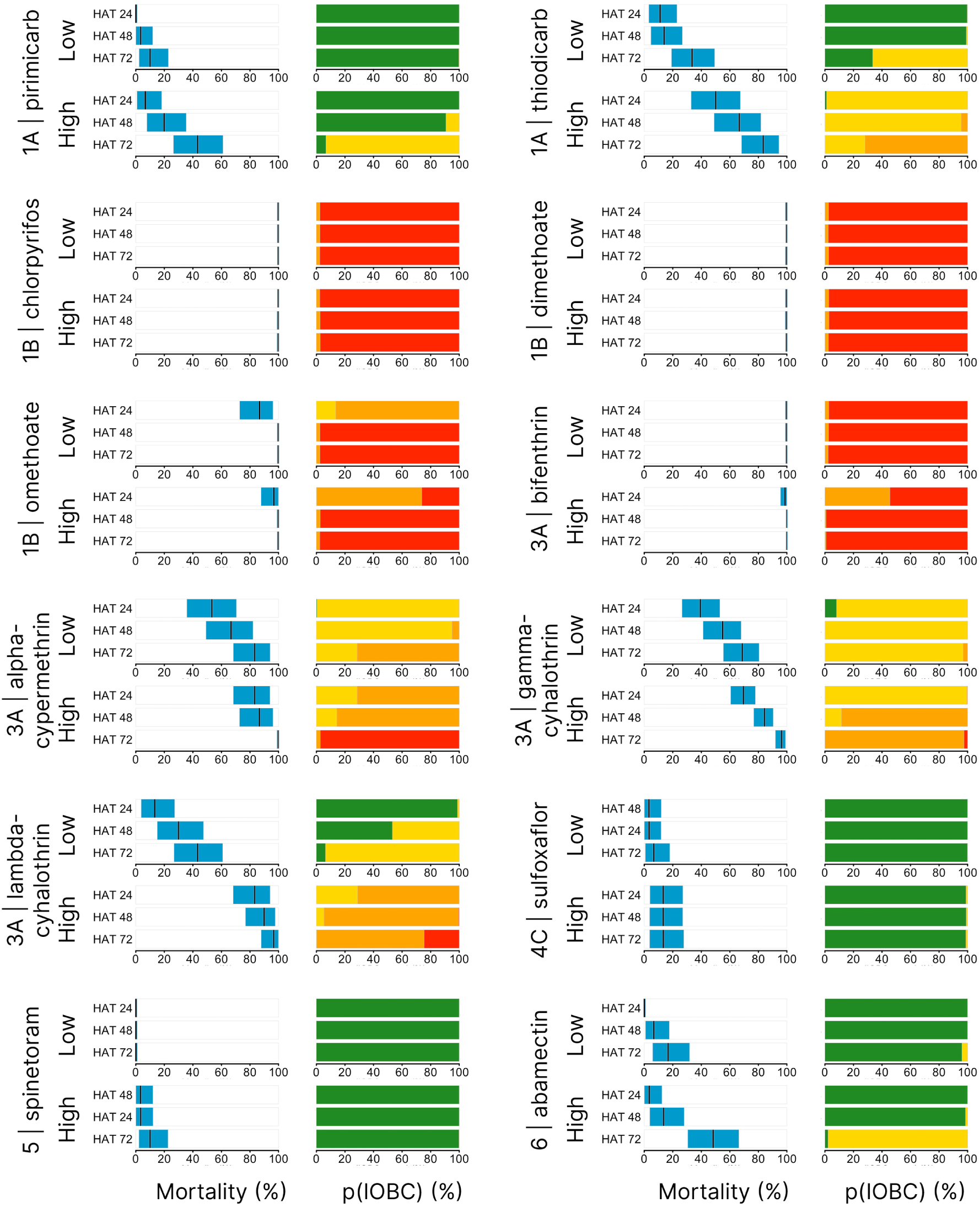
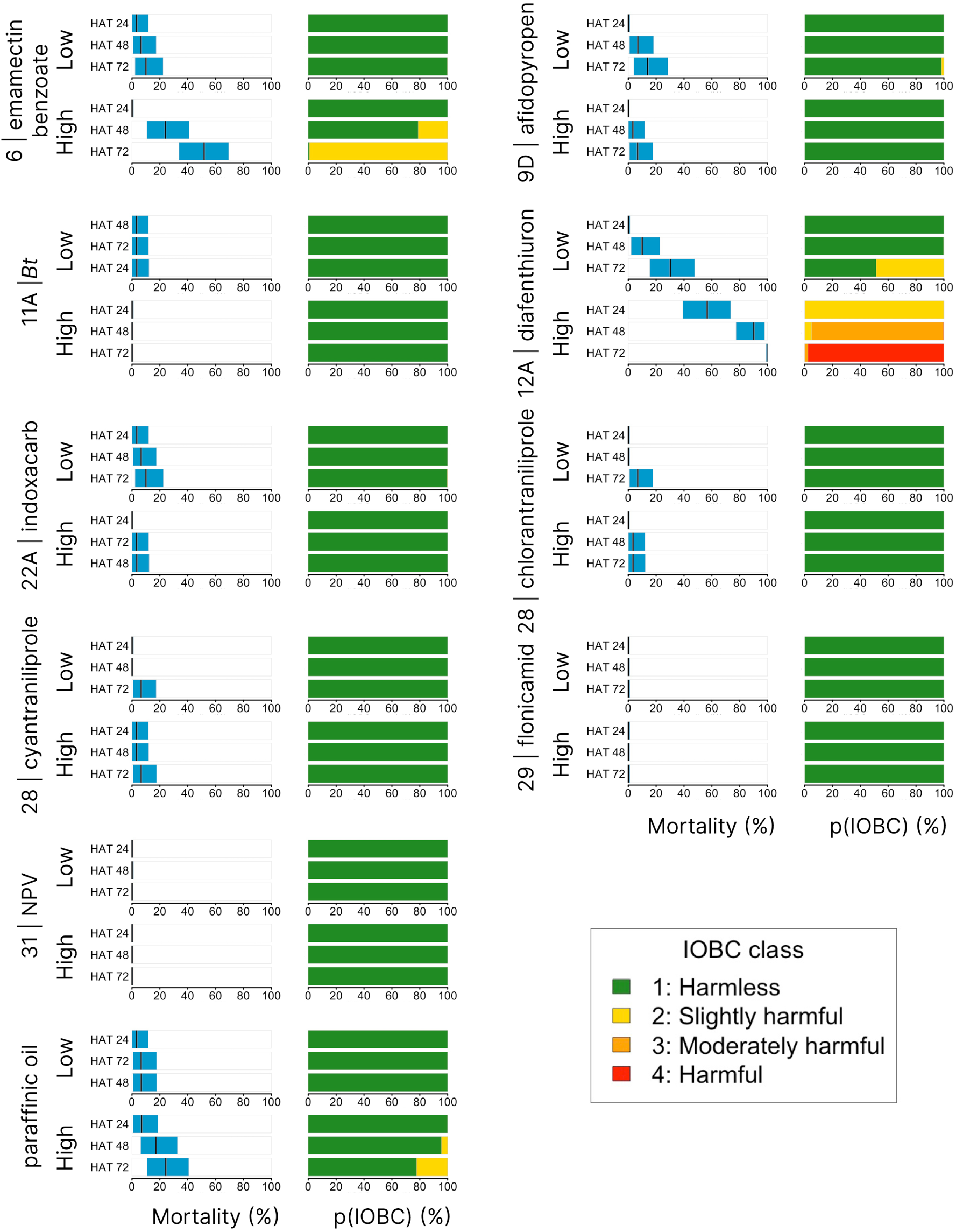
*O. lapidaria* mortality following exposure to pesticide residues at different rates and timepoints. Pesticide rates are described in Table 1. Black lines indicate the mean response and the blue rectangles the associated statistical uncertainty (95% CI). Non-overlapping rectangles denote statistical differences. Pairwise statistical comparisons and mean group effects are given in Table S4. The right-hand panels show the probability or probabilities that the mortality response of *O. lapidaria* to the chemical, rate and HAT treatments will be categorised into one or more of the IOBC toxicity classifications, where 1 = ‘harmless’ (< 30% mortality; green), 2 = ‘slightly harmful’ (30-79% mortality; yellow), 3 = ‘moderately harmful’ (80-99% mortality; orange), and 4 = ‘harmful’ (> 99% mortality, red). p: probability; IOBC: International Organisation for Biological Control; HAT: hours after treatment; CI: credible interval.

Pesticide exposure time was also an important determinant of *O. lapidaria* mortality for some active ingredients. This effect was particularly clear for emamectin benzoate at the high rate, where mortality at 24-HAT to 72-HAT increased from 0.1% to 51.7% (Figure 2; Table 1). This corresponds with a statistically different average increase in toxicity of 47.1-fold (Table S4). The other active ingredients which caused statistical differences in average *O. lapidaria* mortality between 24-HAT and 72-HAT were gamma-cyhalothrin, omethoate and pirimicarb (when tested at low rates), alpha-cypermethrin (when tested at the high rate), and abamectin, afidopyropen and diafenthiuron (at both rates tested)(Figure 2; Table S4). On the other hand, exposure time had no impact on *O. lapidaria* mortality for the rapidly acting ‘harmful’ active ingredients chlorpyrifos, dimethoate, and bifenthrin, each of which caused on average 100% mortality by 24-HAT when tested at high rates (Figure 2; Table 1). There were no statistical differences in *O. lapidaria* mortality between 24-HAT and 72-HAT for these three active ingredients, as well as for *Bt*, NPV, chlorantraniliprole, cyantraniliprole, flonicamid, indoxacarb, lambda-cyhalothrin, paraffinic oil, spinetoram, and sulfoxaflor, at either rate tested (Table S4).

### 3.3. Chemical responses between species (RQ3)

*Odontoscirus lapidaria* and Bdellidae sp. B showed consistent responses to four of the five representative pesticides tested at 48-HAT (Figure 3; Table S5), with no statistical differences in mortality between species for abamectin, paraffinic oil, spinetoram and diafenthiuron at high rates. For both species, abamectin, paraffinic oil and spinetoram were classified as ‘harmless’ (IOBC class 1; <30% mortality), and diafenthiuron was classified as ‘moderately harmful’ (IOBC class 3; 80-99% mortality). However, Bdellidae sp. B was statistically more tolerant to chlorpyrifos than *O. lapidaria*; at 48-HAT mortality was on average five times lower in Bdellidae sp. B (20%) than in *O. lapidaria* (100%). Chlorpyrifos was therefore classified as ‘harmful’ (IOBC class 4; >99%) to *O. lapidaria*, but ‘harmless’ (IOBC class 1; <30% mortality) to Bdellidae sp. B.

**Figure 3 –.**
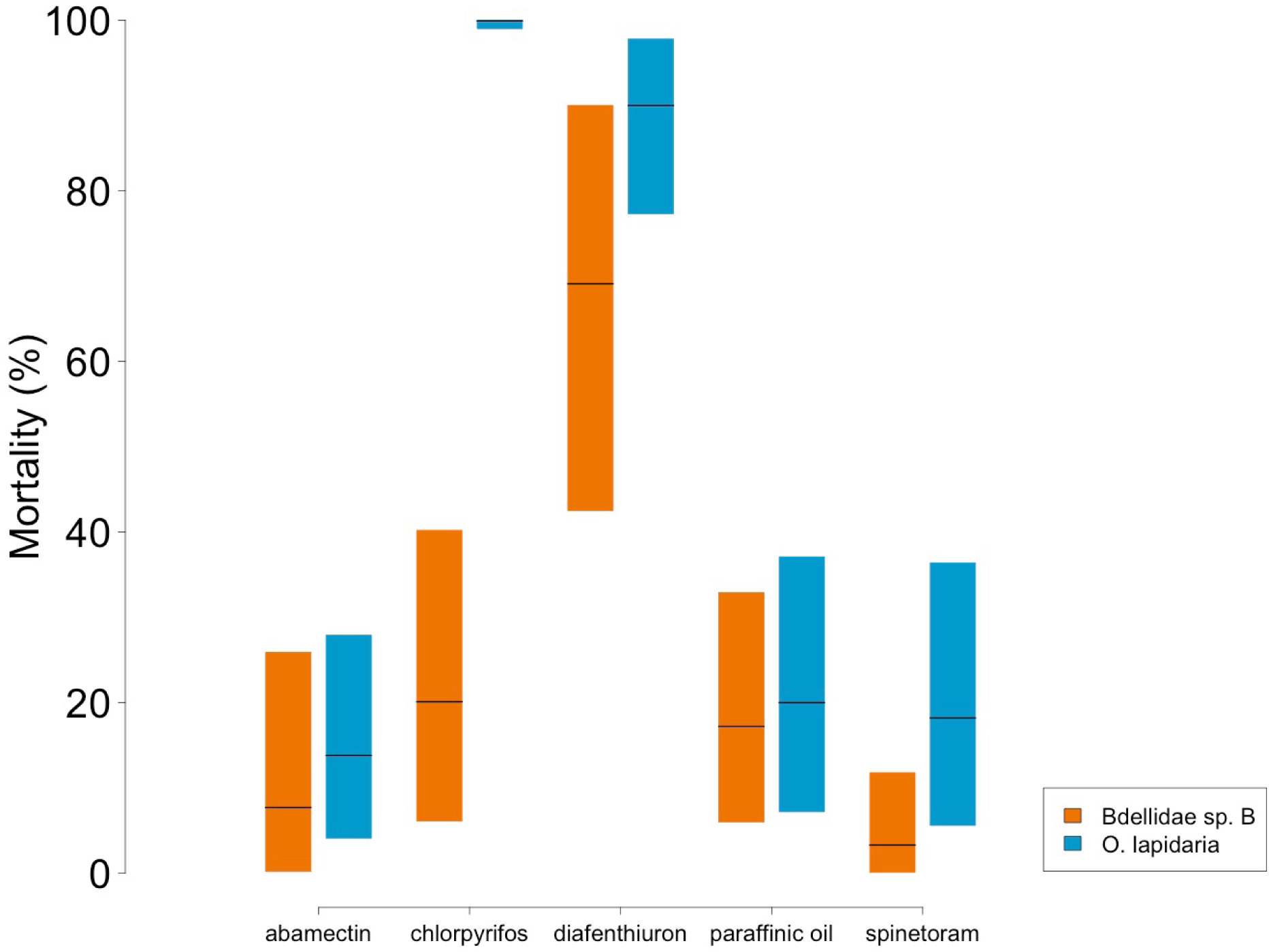
Mortality of *O. lapidaria* and Bdellidae sp. B following exposure to high rates of pesticide residues at 48-HAT. Pesticide rates are described in Table 1. Black lines indicate the mean response and the coloured rectangles the associated statistical uncertainty (95% CI). Non-overlapping rectangles denote statistical differences. HAT: hours after treatment; CI: credible interval.

## 4. Discussion

This study provides standardised pesticide toxicity data for two species of mites from the Bdellidae family in keeping with previous work (e.g. Overton et al., 2023) and standardised IOBC benchmarks (Sterk et al., 1999). To the best of our understanding, this represents the first comprehensive study of pesticide impacts on Bdellidae, with most previously available information on predatory mites from the commercially valuable Phytoseiidae family (Biobest, 2019; Bostanian, 1985, 2006; Choi, 2003; Cote, 2002; Cuthberson, 2012; Duso, 1992; Kaplan et al., 2012; Kim, 2005; Kishimoto, 2018; Zhang, 1990).

Although focused on *O. lapidaria*, we also compared chemical responses with a second Bdellidae species collected from a separate location. While formal identification using available taxonomic keys has not been completed, DNA barcoding confirmed this species (which we refer to as Bdellidae sp. B) is from the Bdellidae family but quite distinct from *O. lapidaria*. The two species showed similar responses to abamectin, diafenthiuron, paraffinic oil and spinetoram. However, important differences in sensitivity to the OP chlorpyrifos were observed, with Bdellidae sp. B showing notably higher tolerance than *O. lapidaria*. This highlights the risk of making generalisations around acute toxicity within natural enemy families, something observed previously for other guilds. For example, Overton et al. (2023) showed the toxicity of the SP gamma-cyhalothrin varies considerably between three parasitoid wasp species important to biological control of crop aphids (*Aphidius colemani*, *Diaeretiella rapae*, *Aphelinus abdominalis*).

In the case of predatory mites, even different populations of the same species may respond differently to the same pesticides (Anber and Overmeer, 1988; Hardman et al., 1991; Hoy and Conley, 1987; Nyrop et al., 1995). This may be due to past selection for resistance (İnak and Yorulmaz, 2022; Kreiter et al., 2010), something which could also contribute to the higher chlorpyrifos tolerance seen in Bdellidae sp. B. The development of pesticide-resistance in some natural enemy species or populations shows promise for IPM, offering a possible resolution to the conflict between biological and chemical pest control (Bielza, 2016). However, it is not yet known whether pesticide exposure influences resistance in different populations of Bdellidae sp. B and in any case this mite is likely to be absent in many fields in Australia (Hyojung Kang et al., unpubl. data).

For many pesticides, acute toxicity responses of *O. lapidaria* were consistent with the responses of Phytoseiidae and other beneficial invertebrates more broadly. Unsurprisingly, *O. lapidaria* showed high sensitivity to broad-spectrum pesticides such as OPs and SPs, in line with past work showing the incompatibility of these chemicals with IPM actions aimed at using predatory mites as biocontrol agents (Agnello et al., 2003). The high toxicity of OPs and SPs towards *O. lapidaria* is largely congruous with previous research on phytoseiid mites (James and Rayner, 1995; Hassan et al., 1988; Bonafos et al., 2007; Miles and Dutton, 2003; Kishimoto et al., 2018) and a range of other beneficial invertebrates (summarised by Knapp et al., 2023B; Overton et al., 2021). Previous work has also demonstrated the harmfulness of SPs and OPs towards bdellids, although these findings have sometimes been conflicting. Roberts et al. (2011) showed that the SPs, bifenthrin and alpha-cypermethrin, and the OPs, chlorpyrifos and omethoate, caused 100% mortality to *O. lapidaria* after only 8 h of exposure to registered field rates, when tested in the laboratory. Similarly, Michael (1991) showed that spray applications of chlorpyrifos in the field significantly reduced population densities of *O. lapidaria*. However, exposure to omethoate across a number of field trials has shown little toxic effect on *O. lapidaria* (Roberts et al., 2011; Michael, 1991, 1995) and *N. capillatus* (family Bdellidae; Michael, 1991), despite causing high mortality in the laboratory (this study and Roberts et al. 2011). This could be due to a number of reasons, including reduced pesticide contact in the field due to vegetation cover or behavioural avoidance (Wallace, 1971), highlighting the challenges of extrapolating laboratory toxicities to field situations.

Our study also revealed several active ingredients that seem to have consistently low impacts on bdellid mites and other beneficial invertebrates more broadly. In keeping with previous work on a range of phytoseiid mites (Bernard et al., 2009; Kim et al., 2011, 2018; Kishimoto et al., 2018) and other beneficial invertebrates (Zantedeschi et al., 2018; Roubos et al., 2014; Angeli et al., 2005; Grundy, 2007), the biological insecticides *Bt* and NPV, which target lepidopteran larvae, were harmless towards *O. lapidaria*. Flonicamid, afidopyropen, cyantraniliprole and chlorantraniliprole also had no observable impact on *O. lapidaria*, consistent with impacts on a diverse array of beneficial species that contribute to biological control, including aphid parasitoids (Overton et al., 2023) and other natural enemies (summarised by Knapp et al., 2023B). Based on our findings, these active ingredients show promise for IPM strategies and are likely to be highly compatible with conservation biological control efforts involving naturally-occurring beneficials such as Bdellidae.

On the other hand, this study has revealed key differences between the impacts of some pesticides on Bdellidae compared with other predatory mite families. Most notably, diafenthiuron is reported to be ‘harmless’ to the phytoseiid species *Amblyseius longispinosus* (Pankaj and Abhishek, 2015), *Amblyseius finlandicus* (Shukla and Mandape, 2017), *T. pyri* and *P. persimilis* (Jansen, 2010), yet in our study this active ingredient caused on average 90% mortality to *O. lapidaria* and 69% mortality to Bdellidae sp. B. In contrast, *O. lapidari*a and Bdellidae sp. B showed tolerance towards abamectin (consistent with previous work on *O. lapidaria*; Roberts et al., 2011), despite this active ingredient being very harmful towards *N. californicus* (family Phytoseiidae; Kaplan et al., 2012). Further research is required to understand the biological factors underpinning variations in pesticide responses between Phytoseiidae and Bdellidae. Differences resulting from body size (Phytoseiidae being 0.2-0.58 mm and Bdellidae reaching up to 3.5 mm; Hernandes et al., 2015; Tixier et al., 2012) and adaptations to different life-histories (Phytoseiidae being leaf-dwelling and Bdellidae being soil-dwelling; Hernandes et al., 2015; Knapp et al., 2018) are likely to contribute. These discrepancies in pesticide tolerance and biology strengthen the case for extending standardised pesticide toxicity testing beyond the commercially available range of species typically studied.

For many active ingredients, the timepoint at which mortality assessments were taken had implications for their IOBC classification. This highlights important nuances to chemical toxicity; while IOBC classifications and methodologies are useful (Sterk et al., 1999), they represent a snapshot of toxicity in one scenario and may over-simplify temporal effects. This is exemplified by other studies; for instance, Lima et al. (2016) examined the long-term impacts of different pesticides on *Neoseiulus baraki* (Phytoseiidae), finding abamectin to cause 100% mortality by eight days after treatment despite having only moderate impacts on mortality initially. In the case of *O. lapidaria*, temporal effects were particularly evident for several active ingredients, including thiodicarb, alpha-cypermethrin, gamma-cyhalothrin, lambda-cyhalothrin, diafenthiuron, abamectin and emamectin benzoate. These active ingredients would particularly benefit from further study across an extended range of rates and timepoints. In addition, the sub-lethal impacts of chemicals not captured by mortality assessments are important to consider alongside IOBC classifications, including metrics such as reproduction, lifespan, learning performance, searching behaviour and neurophysiology (Desneux et al., 2007; Duso et al., 2020). For instance, Bostanian and Akalach (2006) found indoxacarb caused a 26.7% reduction in fertility of *P. persimilis* despite having little impact on mortality.

## 5. Conclusions

Standardised toxicity assessments are an important step in identifying selective pesticides compatible with IPM. This study is the first comprehensive assessment of pesticide impacts on Bdellidae and has illuminated key contrasts within predatory mites. It seems that responses of predatory mites to agricultural pesticides can vary between families, between species within the same family, as well as temporally within the same specie - as demonstrated here and also seen in other studies (e.g. Hoy and Conley, 1987; Jansen, 2010; Pankaj and Abhishek, 2015; Shukla and Mandape, 2017). This highlights the difficulties of making generalisations around pesticide toxicity towards predatory mites. As such, conservation biological control using predatory mites will be best supported when the diversity of species included in standardised toxicity assessments reflects the diversity of naturally-occurring species contributing to biological control.

## CRediT authorship contribution statement

Funding acquisition: AAH, PAU; Conceptualisation: RAK, LM, AAH, PAU; Methodology: RAK, LM, RM, QY, AAH, PAU; Formal analysis: LM; Writing – original draft: RAK; Writing – review and editing: LM, RM, QY, AHH, PAU; Visualisation: RAK, LM; Supervision: LM, PAU; Project administration: RAK, PAU.

## Data availability statement

Data and codes to reproduce models and plots are already published and publicly available in Zenodo: https://doi.org/10.5281/zenodo.8368727 (Knapp et al., 2023A) and https://doi.org/10.5281/zenodo.8368898 (Umina et al., 2023).

## Acknowledgements

This research was funded by the Grains Research and Development Corporation, grant number UOM1906–002RTX. We would like to thank Jade Russell, Ana Giraldo, Alex Slavenko, Tara Jalali, Lisa Kirkland, Adriana Arturi and Melissa Carew for technical assistance. We acknowledge the Traditional Custodians of the land on which our research was conducted and pay our respects to their Elders past and present.

## Supplementary Materials

**Figure S1 –.**
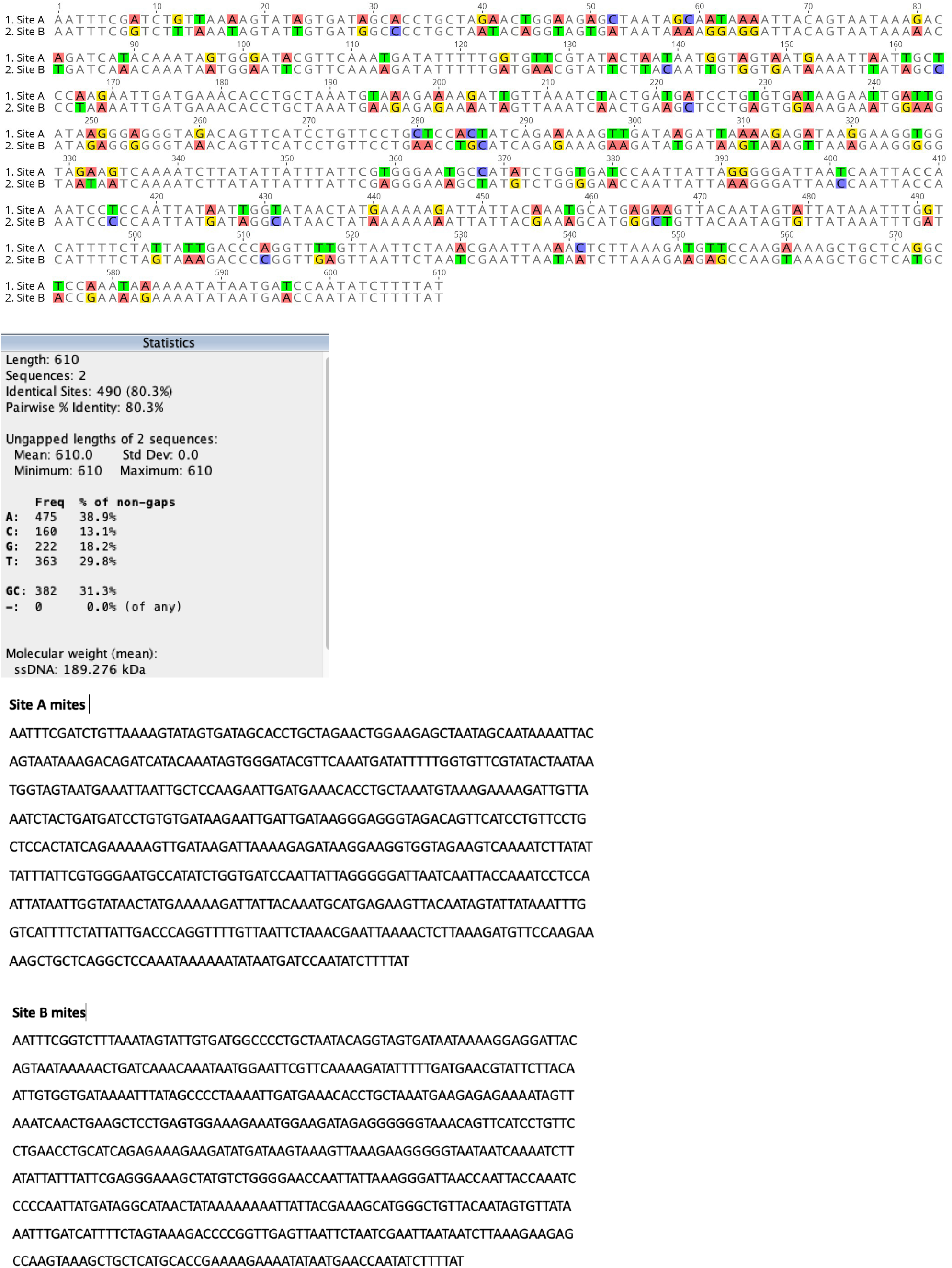
Sequence alignment of Site A mites (***O. lapidaria***) and Site B mites (Bdellidae sp. B).

**Table S1 –** Negative control mortality and chemical treatments tested in each bioassay involving *O. lapidaria* and Bdellidae sp. B. N values below 30 indicate cases where mites escaped, were killed or injured during handling, or where insufficient numbers could be obtained from field collections. The same active ingredient was occasionally tested over multiple bioassays (i.e. thiodicarb, gamma-cyhalothrin and bifenthrin). In these cases, results were pooled across bioassays.

**Table S2 –** Pairwise comparisons for *O. lapidaria* mortality by chemical model, with cell values representing the mean group effect (MGE) as described in the main text. Chemical treatments with overlapping 95% CIs are noted as ‘not statistically different’ (nsd). Means and CIs are given in Table S3.

**Table S3 –** Probability (p) that the mortality response of *O. lapidaria* to a given chemical at a given rate and HAT will be classified into one or more of the IOBC toxicity categories: 1 = ‘harmless’ (<30% mortality); 2 = ‘slightly harmful’ (30-79% mortality); 3 = ‘moderately harmful’ (80-99% mortality); and 4 = ‘harmful’ (>99% mortality). Probabilities were derived from posterior estimate summaries for the mortality by chemical and mortality by chemical and HAT models.

**Table S4 –** Pairwise comparison tables for the *O. lapidaria* mortality by chemical and HAT model at low and high rates, with cell values representing the mean group effect (MGE) as described in the main text. HAT timepoints with overlapping 95% Credible Intervals (CI) are noted as ‘not statistically different’ (nsd). Means and CIs are given in Table S3.

**Table S5 –** Posterior estimate summaries for the mortality by chemical and species models comparing *O. lapidaria* and Bdellidae sp. B.

Tables S1 – S5 are provided in a separate Excel spreadsheet.

